# Ray tracing 3D spectral scenes through human optics models

**DOI:** 10.1101/589234

**Authors:** Trisha Lian, Kevin J. MacKenzie, David H. Brainard, Nicolas P. Cottaris, Brian A. Wandell

**Affiliations:** Department of Electrical Engineering, Stanford University, Palo Alto, CA, USA; Facebook Reality Labs, Redmond, WA, USA; Department of Psychology, University of Pennsylvania, Pennsylvania, PA, USA; Department of Psychology, Stanford University, Palo Alto, CA, USA

**Keywords:** 3d rendering, schematic eye models, physiological human optics, retinal image simulation, cone mosaic

## Abstract

Scientists and engineers have created computations and made measurements that characterize the first steps of seeing. ISETBio software integrates such computations and data into an open-source software package. The initial ISETBio implementations modeled image formation (physiological optics) for planar or distant scenes. The ISET3d software described here extends that implementation, simulating image formation for three-dimensional scenes. The software system relies on a quantitative computer graphics program that ray traces the scene radiance through the physiological optics to the retinal irradiance. We describe and validate the implementation for several model eyes. Then, we use the software to quantify the impact of several physiological optics parameters on three-dimensional image formation. ISET3d is integrated with ISETBio, making it straightforward to convert the retinal irradiance into cone excitations. These methods help the user compute the predictions of optics models for a wide range of spatially-rich three-dimensional scenes. They can also be used to evaluate the impact of nearby visual occlusion, the information available to binocular vision, or the retinal images expected from near-field and augmented reality displays.

## Introduction

Vision is initiated by the light rays entering the pupil from three-dimensional scenes. The cornea and lens (physiological optics) transform these rays to form a two-dimensional spectral irradiance image at the retinal photoreceptor inner segments. How the physiological optics transforms these rays, and how the photoreceptors encode the light, limits certain aspects of visual perception and performance. These limits vary both within an individual over time and across individuals, according to factors such as eye size and shape, pupil size, lens accommodation, and wavelength-dependent optical aberrations (Wyszecki & Stiles, 1982).

The physiological optics is accounted for in vision science and engineering by a diverse array of models. In certain cases the application is limited to central vision and flat (display) screens, and in these cases the optical transformation is approximated using a simple formula: convolution with a wavelength-dependent point spread function (Wandell, 1995). This approximation is valid when the stimulus is paraxial and distant. But in other important cases, for example when viewing nearby 3D objects, the optical transformation is more complex and depends on the three-dimensional geometry and spatial extent of the scene. For example, in the near field, the point spread function is depth dependent, and shift-invariant convolution produces incorrect approximations of retinal irradiance at image points near depth occlusions. Accurately calculating the optical transformation for scenes within 1-2 meters of the eye, which are important for understanding depth perception and vergence, requires more complex formulae and computational power.

This paper describes ISET3d, a set of software tools that simulates the physiological optics transformation from a three-dimensional spectral radiance into a two-dimensional retinal image. The software is integrated with ISETBio, an open-source package that includes a number of computations related to the initial stages of visual encoding (Cottaris, Jiang, Ding, Wandell, & Brainard, 2018; Brainard et al., 2015; Kupers, Carrasco, & Winawer, 2018). The initial ISETBio implementations modeled image formation for planar or distant scenes with wavelength-dependent point spread functions (Farrell, Catrysse, & Wandell, 2012; Farrell, Jiang, Winawer, Brainard, & Wandell, 2014). ISET3d uses quantitative computer graphics to model the depth-dependent effects of the physiological optics and enables the user to implement different schematic eye models for many different three-dimensional scenes. For some of these models the software can calculate retinal irradiance for a range of accommodative states, pupil sizes and retinal eccentricities.

ISETBio uses the retinal irradiance to estimate the excitations in the cone spatial mosaic. These estimated cone excitations can be helpful in understanding the role of the initial stages of vision in limiting visual performance, including accommodation as well as judgments of depth and size. In addition, these calculations can support the design and evaluation of 3D display performance (e.g. light fields, augmented reality, virtual reality), which benefit from a quantitative understanding of how display design parameters impact the retinal image and cone excitations. For example, ISET3d may help develop engineering metrics for novel volumetric displays that render rays as if they arise from a 3D scene, or for multi-planar displays that use elements at multiple depths to approximate a 3D scene (Akeley, Watt, Girshick, & Banks, 2004; MacKenzie, Hoffman, & Watt, 2010; Mercier et al., 2017; Narain et al., 2015). In summary, we describe ISET3d as a tool that can be helpful to various different specialties within vision science and engineering.

## Methods

ISET3d combines two key components: quantitative computer graphics based on ray-tracing and a method for incorporating eye models within the computational path from scene to retinal irradiance (Figure 1). The calculations begin with a 3D graphics file that defines the geometry, surfaces, and lighting of a scene. The rendering software traces rays in the scene through a configurable eye model that is defined by a series of curved surfaces and an aperture (pupil). The rays arrive at the surface of the curved retina, where they form the retinal irradiance (optical image). Using ISETBio, the retinal irradiance can be sampled by a simulated cone photoreceptor mosaic, and an expected spatial pattern of cone excitations can be calculated

**Figure 1:**
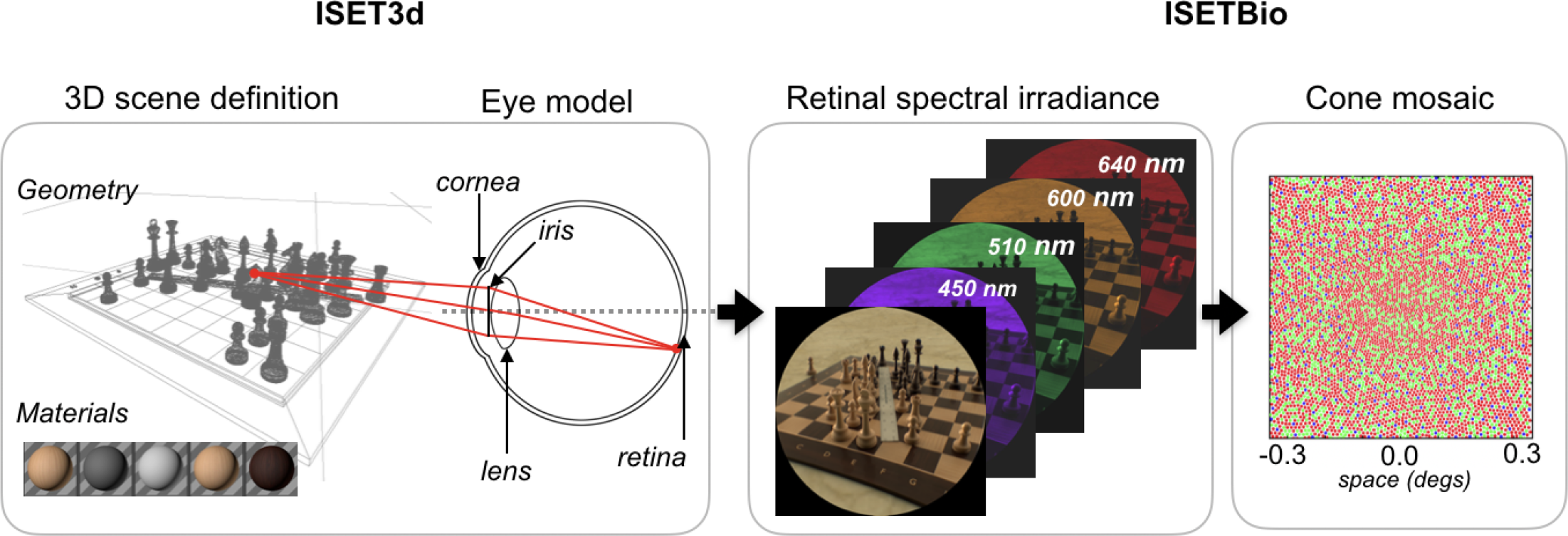
The computational pipeline. A three-dimensional scene, including objects and materials, is defined in the format used by Physically Based Ray Tracing (PBRT) software (Pharr et al., 2016). The rays pass through an eye model implemented as a series of surfaces with wavelength-dependent indices of refraction. The simulated spectral irradiance at the curved retinal surface is calculated in a format that can be read by ISETBio (Cottaris, Jiang, et al., 2018). That software computes cone excitations or current at the cone outer segment membrane, in the presence of fixational eye movements (Cottaris, Rieke, et al., 2018).

### PBRT integration

We implement rendering computations using the open-source PBRT software (Pharr et al., 2016). This ray tracing package calculates physically accurate irradiance (renderings) of 3D scenes. The software incorporates camera models that are defined by multi-element lenses and apertures. We extend the PBRT camera models to implement eye models. These are specified by surface position, curvature, and wavelength-dependent index of refraction; aperture position and size; and retinal position and curvature. The eye model in the ISET3d implementation is sufficiently general to incorporate a variety of physiological eye models.

The software also includes methods to help the user create and programmatically control PBRT scene files that are imported from 3D modeling software (e.g. Blender, Maya, Cinema4D). We implemented ISET3d functions to read and parse the PBRT scene files so that the user can programmatically control scene parameters, including the object positions and orientations, lighting parameters, and eye position (Lian, Farrell, & Wandell, 2018). The scenes used in this paper can be read directly into ISET3d.

The PBRT software and these physiological optics extensions are complex and include many library dependencies. To simplify use of the software, we created a Docker container^1^ that includes the software and its dependencies; in this way users can run the software on most platforms without further compilation. To run the software described here the user installs Docker and the ISET3d and ISETBio Matlab toolboxes.

### PBRT modifications

Ray tracing computations typically cast rays from sample points on the image plane towards the scene; in this application the image plane is the curved inner segment layer of the retina. Rays from the retina are directed towards the posterior surface of the lens. As they pass through the surfaces and surrounding medium of the physiological optics, they are refracted based on Snell’s law and the angle and position of each surface intersection (*Snell’s Law*, 2003). To model these optical effects we extended the main distribution of PBRT in several ways.

First, the original lens tracing implementation was modified to account for the specific surface properties of physiological eye models (e.g. aspheric, biconic surfaces). Second, each traced ray is assigned a wavelength, which enables the calculation to account for the wavelength-dependent index of refraction of each ocular medium. Once the ray exits the physiological optics and enters the scene, standard path tracing techniques within PBRT calculate the interaction between the ray, the material properties of the scene assets, and the lighting.

The modified PBRT calculation returns a multispectral retinal irradiance in relative units. To convert to absolute physical units we set the spatial mean of the retinal irradiance equal to 1 lux times the pupil area. This level may be adjusted for other analyses, such as modeling scenes with different illumination levels. There are additional methods for bringing ray-traced scenes into calibrated units (Heasly, Cottaris, Lichtman, Xiao, & Brainard, 2014).

### Lens transmittance

The transmittance of the human lens varies strongly with wavelength (Stockman, 2001). As a result, there is a striking color difference between the rendering of the scene radiance and the retinal irradiance (Figure 2). ISET3d applies the lens transmittance when calculating the retinal irradiance. The assumed lens density is a parameter that can be set. The macular pigment, the other major inert pigment, is not included in the ISET3d representation. Rather, the macular pigment is part of the ISETBio calculation that converts the retinal irradiance into the cone excitations.

**Figure 2:**
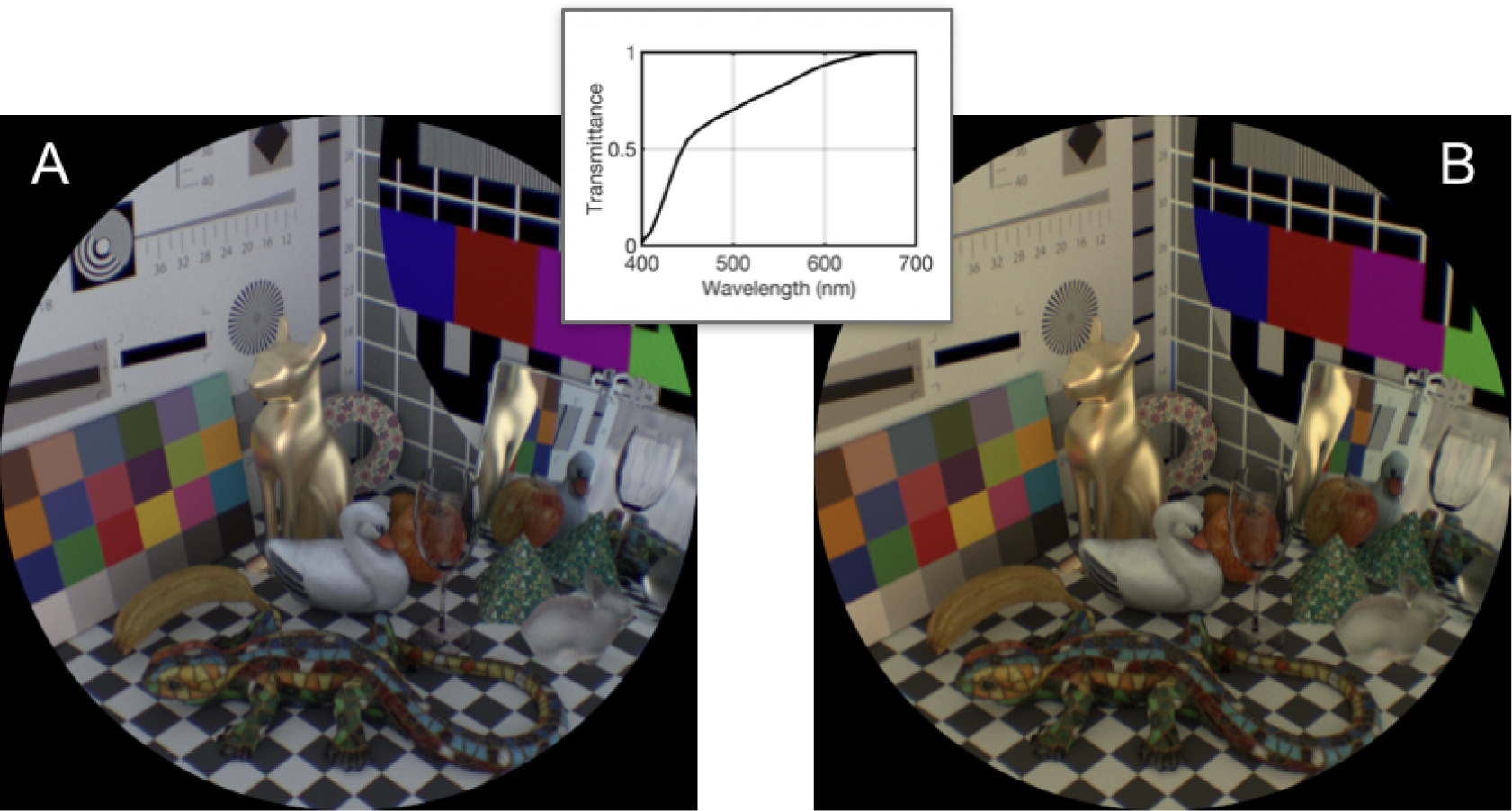
Lens transmittance. The two rendered images illustrate the impact of lens transmittance on the rendered image. Without including lens transmittance (A) the images are relatively neutral in color; including lens transmittance reduces the short-wavelengths and images have a yellow cast (B). The inset at the top is the standard lens transmittance for a 20 year-old.

### Diffraction

Diffraction is a significant contributor to physiological optics blur for in-focus points when the pupil diameter is 2 millimeters (Wandell, 1995). When using computer graphics software to ray-trace or render an image, the effects of diffraction are not typically included. Hence, we added a diffraction module to the PBRT code, based on the technique of Heisenberg Uncertainty Ray-Bending (HURB) (Freniere, Groot Gregory, & Hassler, 1999). HURB is a stochastic model: as each ray passes through the limiting aperture of the optical system the algorithm randomizes the ray direction. The randomization function is a bivariate Gaussian in which the main variance (major axis) is oriented in the direction of the closest aperture point. The sizes of the variance terms depend on the distance between the aperture and the ray, as well as the ray’s wavelength. The degree of randomization approximates optical blur due to diffraction. As the pupil size decreases, relatively more rays come near the edge of the aperture and are scattered at larger angles, thus increasing rendered optical blur.

## Results

First, we validate the ISET3d implementation by comparing the computed modulation transfer function of a schematic eye with calculations and measurements from other sources. Second, we show images modeled using three different eye models. Third, we illustrate the impact of pupil size, accommodation, longitudinal chromatic aberration, and transverse chromatic aberration on the retinal image. Finally, we incorporate ISETBio into the calculation to show how image formation impacts foveal cone excitations.

### Validation

To validate the ISET3d computations, we compare the on-axis, polychromatic modulation transfer function (MTF) of the Navarro schematic eye (Escudero-Sanz & Navarro, 1999) with the MTF computed using Zemax. The ISET3d MTF is derived by ray tracing a slanted-edge, one side black and the other side white, through the Navarro eye model. The irradiance data of the rendered edge are analyzed at each wavelength to calculate the MTF (Williams & Burns, 2014). The wavelength-specific MTFs are combined across wavelength using the photopic luminosity function weighting. The MTFs calculated using ISET3d and Zemax are very similar (Figure 3).

**Figure 3:**
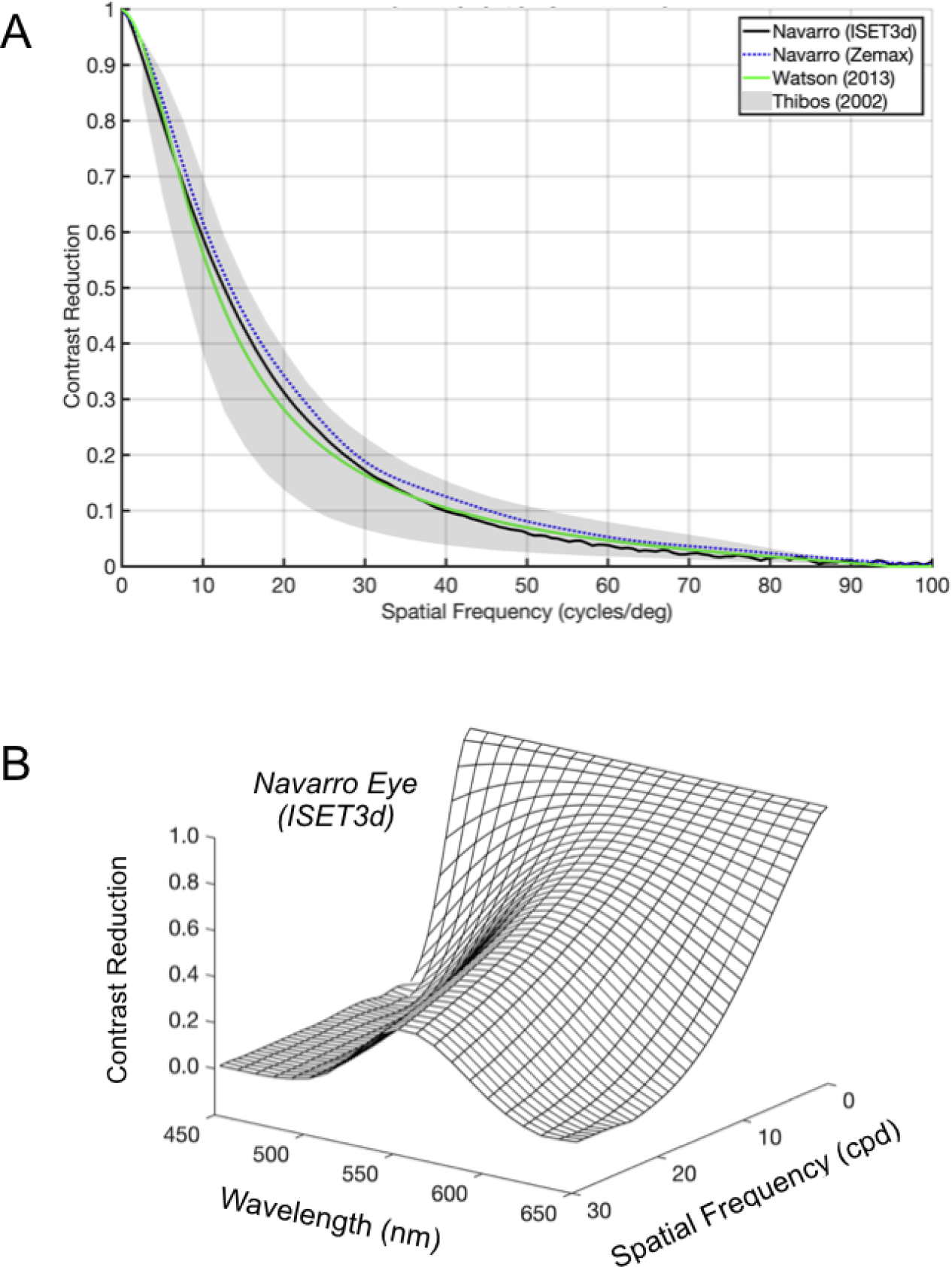
On-axis modulation transfer functions (MTF) of the human optics (pupil diameter 3 mm). (A) A comparison of theoretical and empirical measurements. Two curves were calculated for the Navarro eye model using ISET3d and Zemax. The third curve is based on a formula derived by (Watson, 2013). The gray shaded region is the range of estimates derived from on-axis adaptive optics measurements (Thibos et al., 2002). The spectral radiance of the simulated stimulus is an equal energy polychromatic light and the curves from different wavelengths were combined using a photopic luminosity function weighting. The differences between the curves from ISET3d, Zemax and Watson are smaller than the measured individual variation in optical MTFs. (B) The MTFs for the Navarro eye, calculated using ISET3d with diffraction (see Methods), are shown separately for different wavelengths.

The polychromatic on-axis MTF also closely matches an analytical formulation of the mean, radial MTF derived from empirical measurements (Watson, 2013). The analytical formula is based on wavefront aberration data collected from 200 eyes and is a function of pupil diameter. To assess whether the small differences between ISET3d, Zemax and the analytical formula are meaningful, we compare these curves with the range of MTF functions measured for 100 randomly generated virtual eyes using a statistical model of the wavefront aberration function (Thibos et al., 2002). The ISET3d and Zemax calculations are close to the derived formula. All three are well within the measured population variation.

The luminosity-weighted MTF is a partial description of the human linespread function in three important ways. First, it fails to make explicit the wavelength-dependence of the MTF which is quite strong (Figure 3B). Second, the MTF is shown for a circularly symmetric optical system; effects of astigmatism are omitted. Third, the MTF expresses the contrast reduction for every spatial frequency but omits frequency-dependent phase shifts. In model eyes these phase shifts are often assumed to be zero, but in reality the frequency-dependent phase shifts are substantial. These phase shifts can be modeled by implementing more complex surface shapes in the eye models.

### Eye model comparisons

There are two approaches to defining schematic eye models. Anatomical models define a correspondence between model surfaces and the main surfaces of the eyes optics, such as the anterior or posterior surface of the cornea and lens, and the model surface properties are set to match experimental measurements (Polans, Jaeken, McNabb, Artal, & Izatt, 2015). For example, the Gullstrand No. 1 exact eye models the lens as a series of nested, homogenous shells with different refractive indices (Gullstrand, 1909; Atchison & Thibos, 2016). Alternatively, phenomenological models eschew a correspondence between anatomy and model surfaces. Instead, these models are designed to approximate the functional properties of image formation. For example, reduced or simplified eyes such as the Gullstrand-Elmsley eye have fewer surfaces than the human eye (Elmsley, 1936).

Anatomical and phenomenological models are both parameterized by the surface positions, shape, thickness, and wavelength-dependent refractive index of a series of surfaces. The variation of refractive index with wavelength is called the dispersion curve and its approximate slope is called the Abbe number; this wavelength-dependent refractive index models longitudinal chromatic aberration (LCA), which is the largest of the eyes aberrations (Wandell, 1995; Wyszecki & Stiles, 1982).

Eye models have been developed and analyzed using powerful optics analysis and design software such as Zemax. Model predictions can be compared with human physiological optics for many different measures (Bakaraju, Ehrmann, Papas, & Ho, 2008). These include comparing the Zernike polynomial coefficients that represent the wavefront aberrations, the modulation transfer function, or the wavelength-dependent point spread function. Some models seek to match performance in paraxial vision and others seek to account for a larger range of field heights. Because of their emphasis on characterizing the optics, such packages have limited image formation capabilities, typically restricting their analyses to points or 2D images.

The ISET3d implementation builds upon these 2D measures by inserting the eye model into the 3D PBRT calculations; this enables us to calculate the impact of the eye model on relatively complicated three-dimensional scene radiances. ISET3d models the physiological optics as a series of curved surfaces with wavelength-dependent indices of refraction, a pupil plane, and a specification of the size and shape of the eye. At present the implementation specifies surface positions, sizes, and spherical or biconic surface shapes. These parameters are sufficient to calculate predictions for multiple eye models. The parameters for three model eyes are listed in Appendix 1: the Arizona eye (Schwiegerling, 2004), the Navarro eye (Escudero-Sanz & Navarro, 1999), and the Le Grand eye (El Hage & Le Grand, 1980; Atchison, 2017). Figure 4 shows a scene rendered through each of these models, and Figure 5 shows the on-axis, polychromatic MTF calculated using ISET3d, for each model eye at 3 mm and 4 mm pupil diameters.

**Figure 4:**
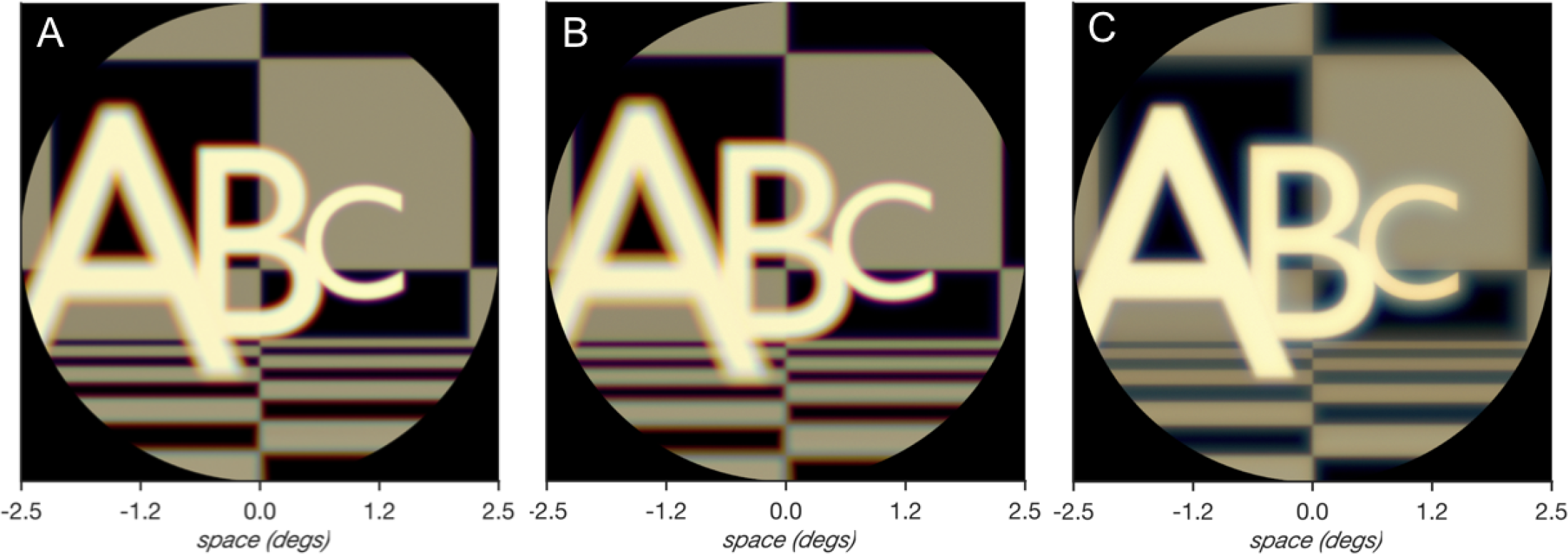
Retinal irradiance calculated using three schematic eye models. (A) Arizona eye (Schwiegerling, 2004) (B) Navarro (Escudero-Sanz & Navarro, 1999) (C) Le Grand (El Hage & Le Grand, 1980). The letters are placed at 1.4 (.714), 1.0 (1), and 0.6 (1.667) diopters (meters) from the eye. The eye models are focused at infinity with a pupil size of 4 mm. Variations in the sharpness of the three letters illustrate the overall sharpness and the depth of field. The images are renderings of the spectral irradiance into sRGB format.

**Figure 5:**
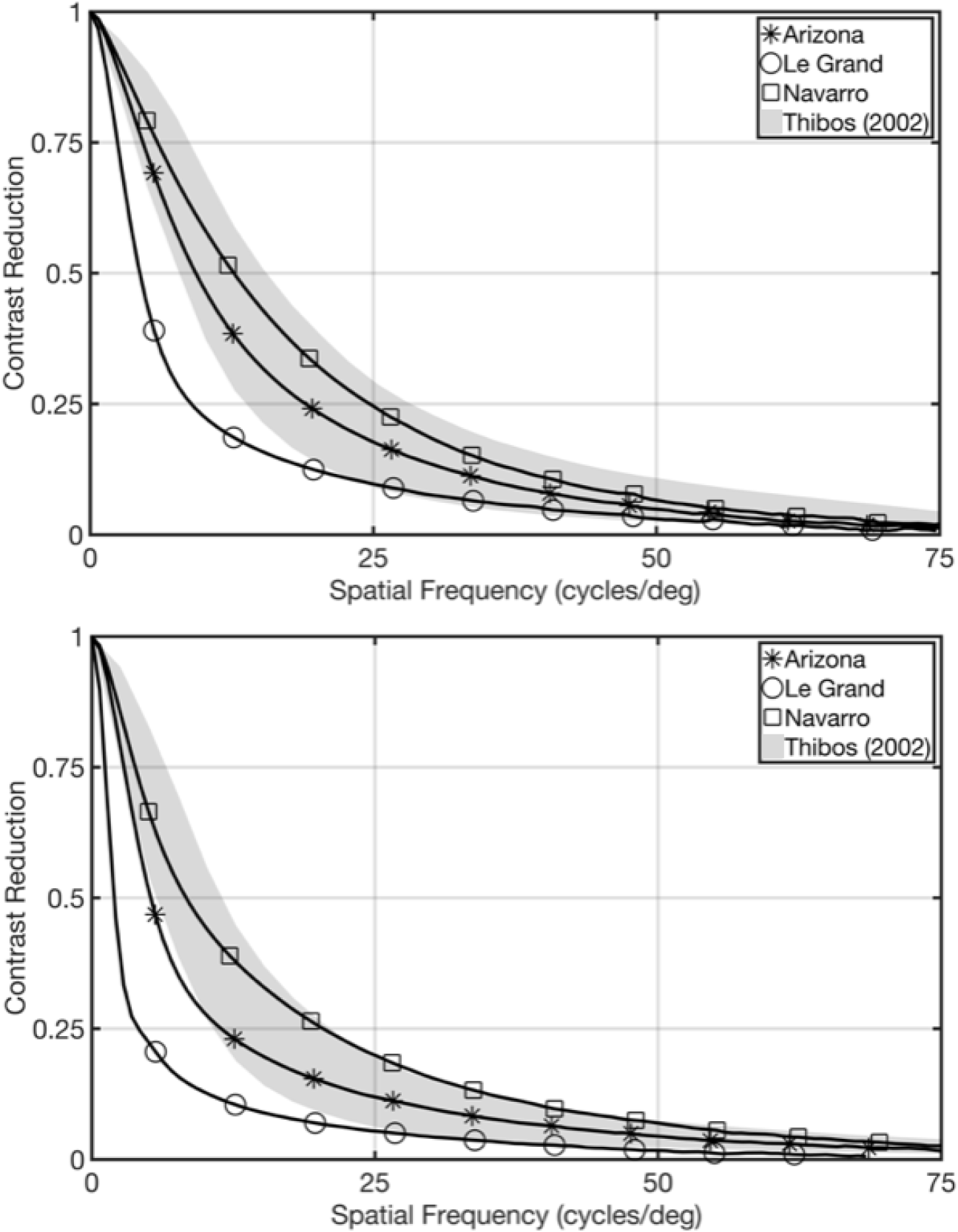
A comparison of on-axis, polychromatic MTF from three different model eyes, calculated using ISET3d. The top figure corresponds to a 3 mm pupil diameter, the bottom figure corresponds to a 4 mm pupil diameter. The gray shaded region is the range of estimates derived from on-axis adaptive optics measurements (Thibos et al., 2002). The Arizona and Navarro eye models are mostly within the range of the measurements; the Le Grand eye is outside the range

For many analyses in this paper, we use the Navarro eye model, although the same calculations can be repeated with any model eye that can be described by the set of parameters implemented in the ray-tracing. The selection of model might depend on the application; for example, analysis of a wide field of view display might require a more computationally demanding model that performs accurately at wide-angles. The goal of this paper is not to recommend a particular eye model, but to provide software tools to help investigators implement, design, evaluate, and use eye models.

### Pupil Size: MTF

The optical performance of the eye changes as the pupil size changes. We measure the change by calculating the MTF using a simulation of the slanted-edge test (Williams & Burns, 2014). The MTF is measured on-axis, and the MTF is calculated for an equal energy wavelength spectral power distribution, with the final result weighted across wavelengths by the luminosity function, as is done in previous sections. The values shown in Figure 6 for the Navarro eye model are qualitatively consistent with the early measurements of the effect of pupil aperture on the width of the line spread function (Campbell & Gubisch, 1966; Wandell, 1995).

**Figure 6:**
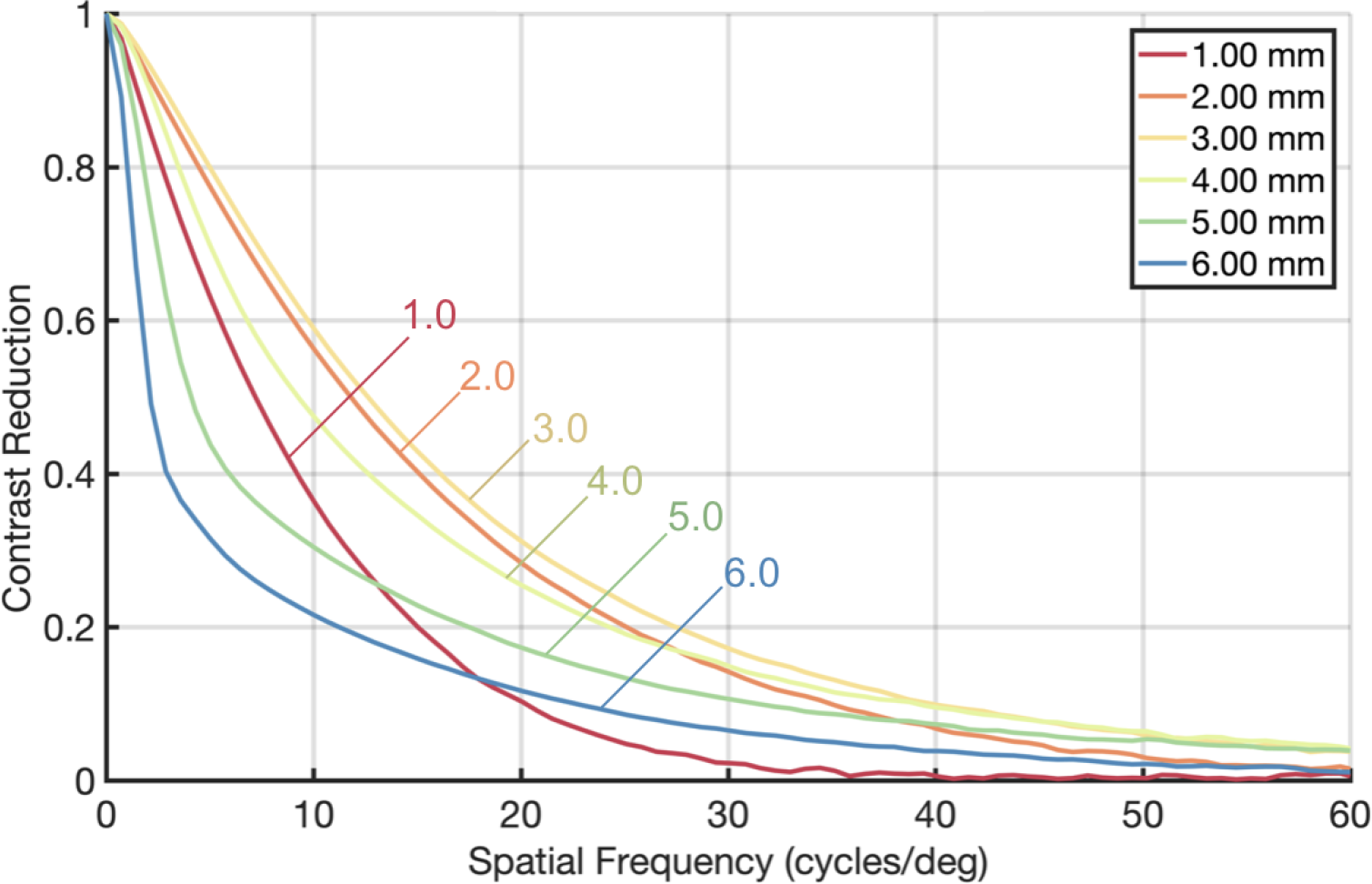
The variation of a model eye modulation transfer function (MTF) with pupil diameter. The on-axis MTF was computed using the Navarro eye model. The best performance is for a 3 mm pupil diameter. The smallest natural pupil size is about 2 mm (Watson & Yellott, 2012). The simulations include the HURB calculation (see Methods) and show that the irradiance from a 1 mm artificial pupil will be impacted by diffraction. The MTF is much lower quality for a large pupil diameter (6 mm).

Decreasing the pupil diameter from 6 mm increases the optical quality up to about 2.5 mm. As the pupil diameter decreases below 2 mm a large fraction of the rays pass near the pupil edge. In this range the scattering effects from the HURB model of diffraction dominate, decreasing the image quality. The HURB model approximates the central part of the diffraction effect (Airy disk), but it does not extend to the first ring.

### Pupil Size: Depth of field

The sharpness of an object’s rendered edge depends on the distance between the object and the accommodation plane. The depth of field is a qualitative property that indicates the range of distances over which objects are in relatively good focus. The smaller the pupil diameter, the larger the depth of field, and the depth of field is much reduced for large (6 mm) compared to small (2 mm) pupil diameters. Figure 7 shows the depth of field for three different pupil diameters, as the eye remains accommodated to a fixed distance.

**Figure 7:**
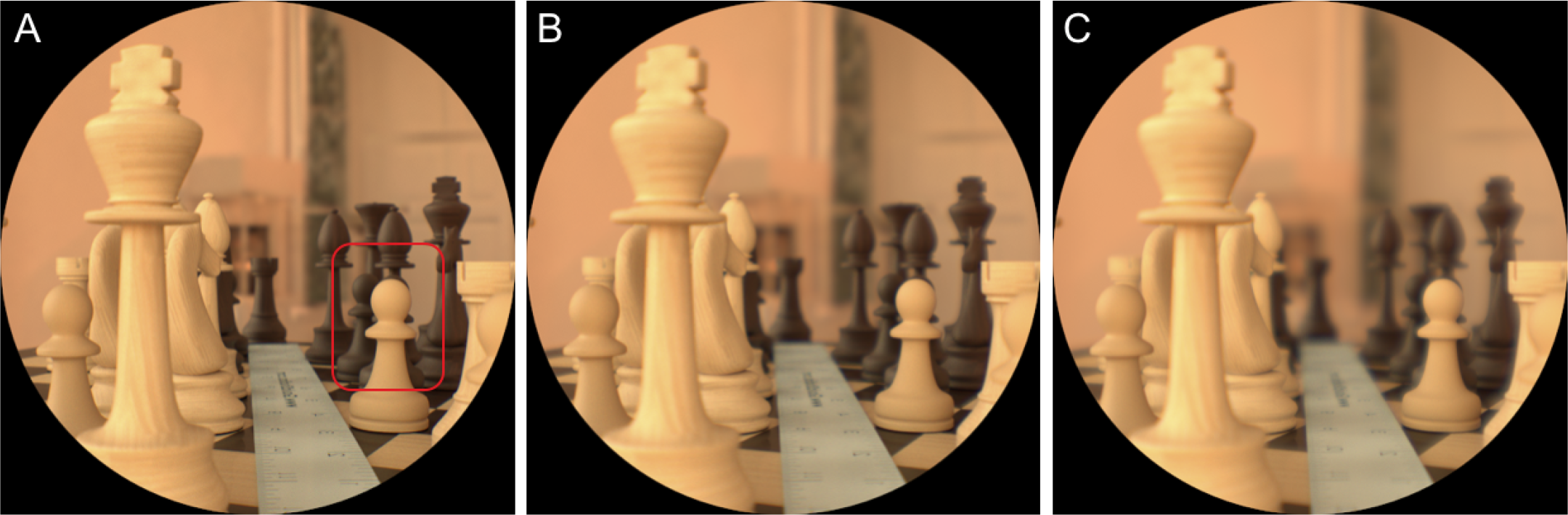
Variations in depth of field calculated for the Navarro eye model with different pupil diameters. Pupil diameters are (A) 2 mm (B) 4 mm, and (C) 6 mm. In all cases, the focus is set to a plane at 3.5 diopters (28 cm), the depth of the pawn shown in the red box. The pawn remains in sharp focus while the chess pieces in front and behind are out of focus; the depth of field decreases as pupil size increases. The horizontal field of view is 30 deg.

### Accommodation

Schematic eyes model accommodation by changing the lens curvature, thickness, and index of refraction according to the formulae provided by the models (e.g. Navarro or Arizona). These parameters can be simulated in ISET3d; hence, we can compute the effect of accommodation on the retinal irradiance. The impact of accommodation using the Navarro eye with a 4 mm pupil diameter is shown in Figure 8. The focus and defocus for the different numbers, presented at 100, 200 and 300 mm distance from the eye, changes substantially as the accommodation of the model eye changes.

**Figure 8:**
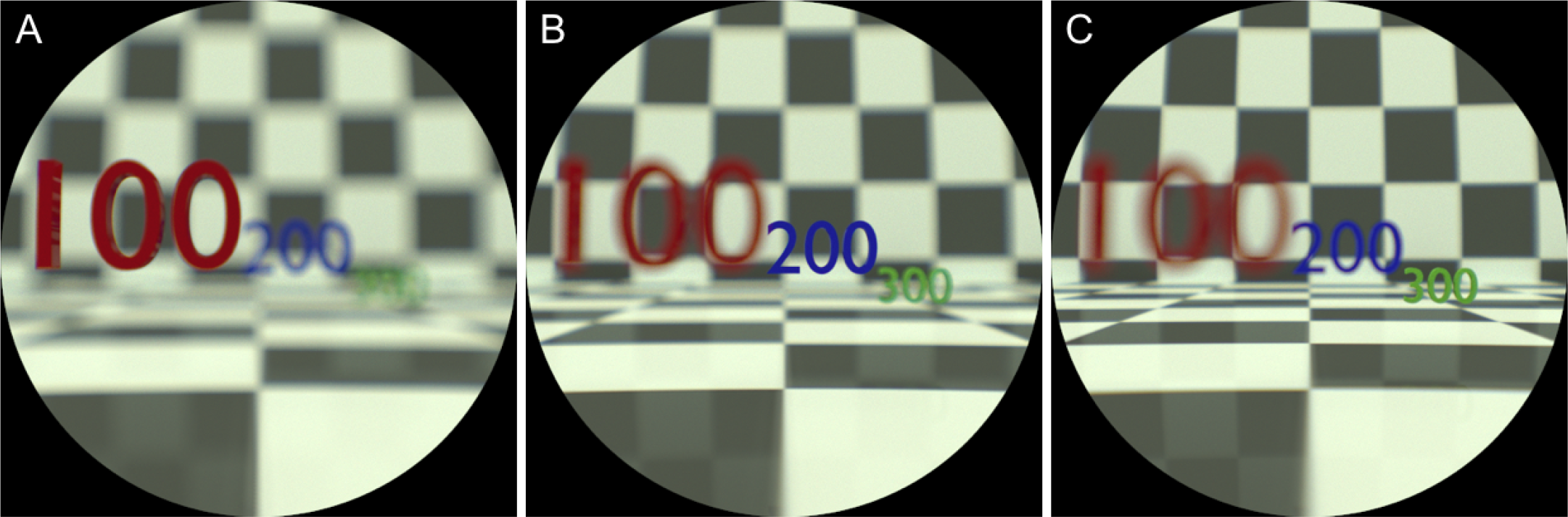
Retinal images for the Navarro eye model accommodated to three target distances. (A) 100 mm, (B) 200 mm, and (C) 300 mm. The images are calculated using a 4 mm pupil diameter. The horizontal field of view is 30 deg.

### Longitudinal Chromatic Aberration

The longitudinal chromatic aberration (LCA) at the eye’s focal plane has been measured several times (Wald & Griffin, 1947; Bedford & Wyszecki, 1957; Wyszecki & Stiles, 1982). The conversion from the LCA measurements (in diopters) to the wavelength-dependent linespread in the focal plane has been worked out (Marimont & Wandell, 1994), and the calculation is implemented in ISETBio for optical models that employ shift-invariant wavelength-dependent point spread function.

ISET3d extends the planar LCA calculation to account for depth-dependent effects. The color fringes at high contrast edges depend on their distance from the focal plane (Figure 9, middle column). The spread of the wavelengths near an edge varies as the eye accommodates to different depth planes. The wavelength-dependent spread at an edge in the focal plane is large for short wavelengths and moderate for long-wavelengths (middle). Accommodating to a more distant plane changes the color fringe at the same edge to red/cyan (top); accommodating to a closer plane changes the chromatic fringe at the edge to blue/orange (bottom).

**Figure 9:**
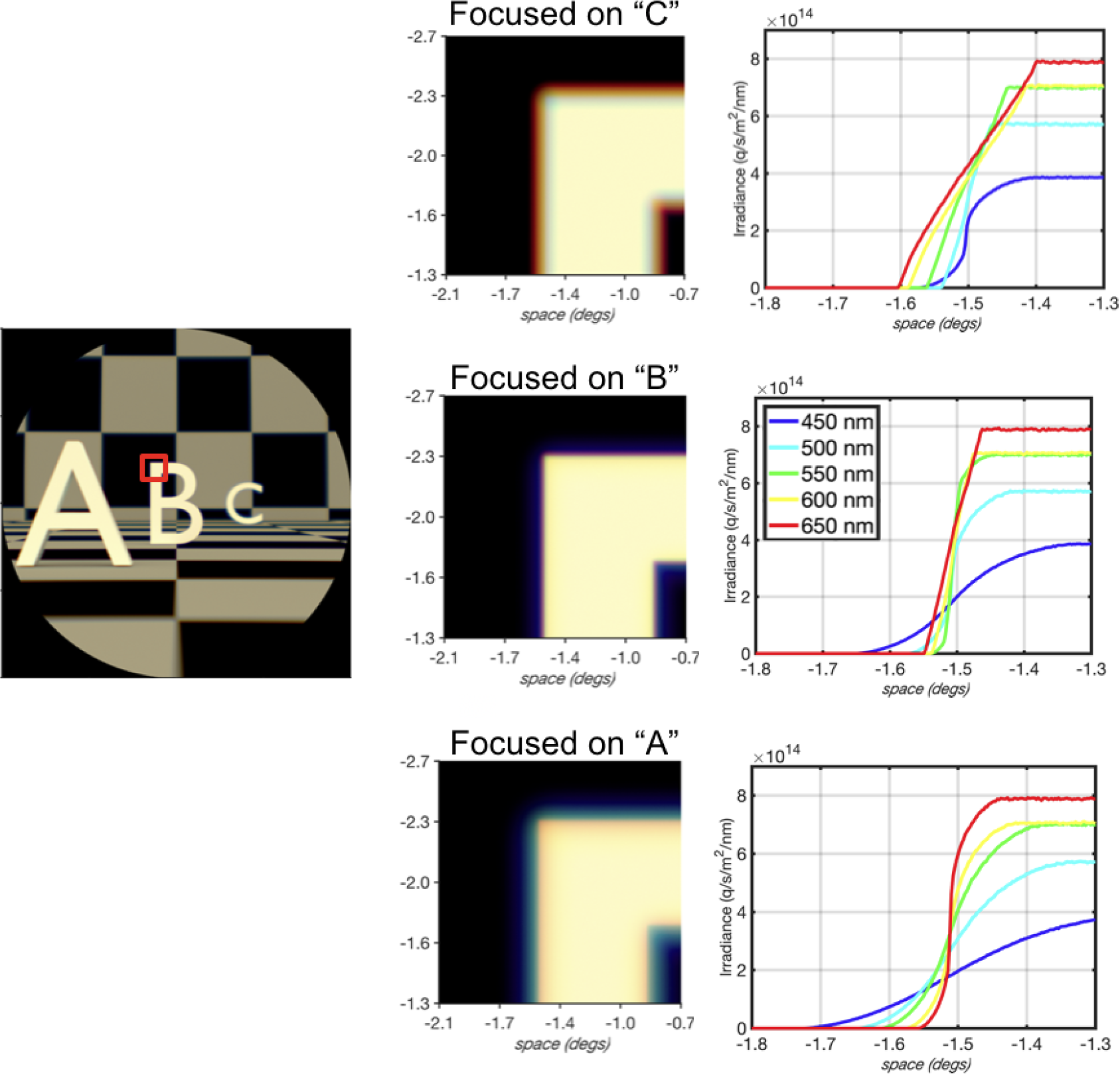
Longitudinal chromatic aberration. A scene including three letters at 1.8, 1.2, and 0.6 diopters (0.56, 0.83, 1.67 m) is the input (left). The scene is rendered three times through the Navarro model eye (4 mm pupil) to form a retinal image with the accommodation set to the different letter depths. The chromatic aberration at the 0.83 m (letter B) depth plane is rendered, showing how the color fringing changes as the focal plane is varied. The graphs at the right show the spectral irradiance across the edge of the target for several different wavelengths.

In these examples the middle-wavelengths spread somewhere between 5-20 minutes of arc; the short-wavelength light spreads over a larger range, from 10-40 minutes of arc. This spread is large enough to be resolved by the cone mosaic near the central retina, and the information in a single image is sufficient to guide the direction of accommodation needed to bring the front or back edge into focus. Experimental results confirm that experimentally manipulating such fringes does drive accommodation in the human visual system (Cholewiak, Love, Srinivasan, Ng, & Banks, 2017).

### Transverse Chromatic Aberration

Transverse chromatic aberration (TCA) characterizes the wavelength-dependent magnification of the image (Thibos, Bradley, Still, Zhang, & Howarth, 1990). TCA arises from several optical factors, including the wavelength dependent refraction of the surfaces and the geometric relationship between the pupil position, scene point, and optical defocus. In any small region of the image, the TCA magnification appears as a spatial displacement between the wavelength components of the irradiance; because the TCA is a magnification, the displacement size increases with eccentricity.

In the human eye the wavelength-dependent shift between 430 nm and 770 nm is approximately 6 arcmin at 15 deg eccentricity (Winter et al., 2016). This shift is larger than the vernier acuity threshold of about 1-2 arcmin at the same eccentricity (Whitaker, Rovamo, MacVeigh, & Mäkelä, 1992). While not massive, the TCA displacement is large enough to be perceived in the periphery (Thibos et al., 1990; Newton, 1984).

Ray tracing through a schematic eye simulates the effect of TCA. This effect can be clearly seen when we calculate the retinal image of a large rectangular grid comprised of white lines on a black background for the Navarro eye model (Figure 10A). The rectangular grid is distorted due to the curvature of the retina. The wavelength dependent magnification (TCA) is evident in the local displacements of different wavelengths. We calculate the shift size between short and long-wavelengths at 15 deg eccentricity to be approximately 10 arcmin, only slightly larger than the measured TCA.

**Figure 10:**
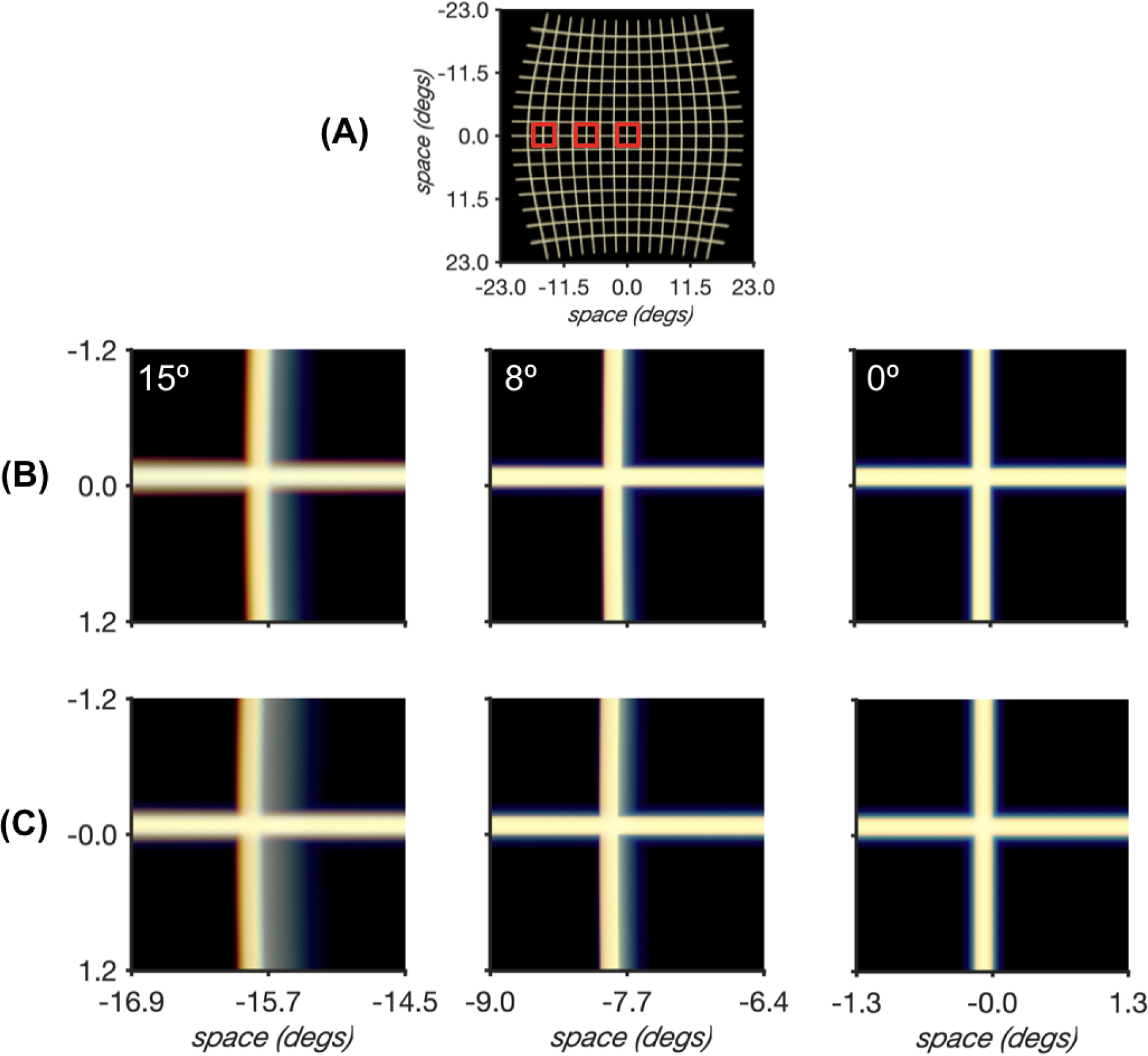
Transverse chromatic aberration (TCA) at different eccentricities and pupil positions. (A) A white grid 1 meter distant is rendered through the Navarro eye, which is accommodated to the grid plane. The curvature of the retina is seen in the distortion of the grid. (B,C) The images in each row show the TCA at 0, 8 and 15 eccentricity. The two rows were calculated after modifying the anterior chamber depth (ACD) of the Navarro eye within anatomical limits (Rabsilber et al., 2006; Boehm et al., 2018). (B) The ACD is set to 3.29 mm. (C) The ACD is set to 2.57 mm. The TCA is larger when the iris and lens are closer to the posterior corneal surface (ACD is smaller).

### Cone mosaic excitations

The ISET3d irradiance calculations can be used as input to the ISETBio methods. These methods transform the retinal irradiance into cone excitations. ISETBio can simulate a wide range of cone mosaic parameters including (a) the hexagonal spacing, (b) the size of an S-cone free zone in the central fovea, (c) the variation in cone spacing with eccentricity (d) the variation in inner segment aperture size and outer segment length with eccentricity, (e) the macular pigment density, (f) the cone photopigment density, and (g) eye movement patterns.

The rate of cone excitations for a briefly presented light-dark edge at the fovea are shown in Figure 11. The overall cone density and aperture sizes change significantly as one calculates from the central foveal to just a few tenths of a degree in the periphery. The exclusion of the S-cones in the central fovea is also quite striking, as is the very low the absorption rates by the S-cones, shown by the relatively large black dots that represent the S-cones (Figure 11D). Their absorption rate is low in large part because the lens absorbs much of the short-wavelength light. Were the S-cone apertures as small as the apertures in the very central fovea, they would receive very few photons. Not shown are the effects of small eye movements, another feature of the visual system that may be simulated by the ISETBio code (Cottaris, Jiang, et al., 2018).

**Figure 11:**
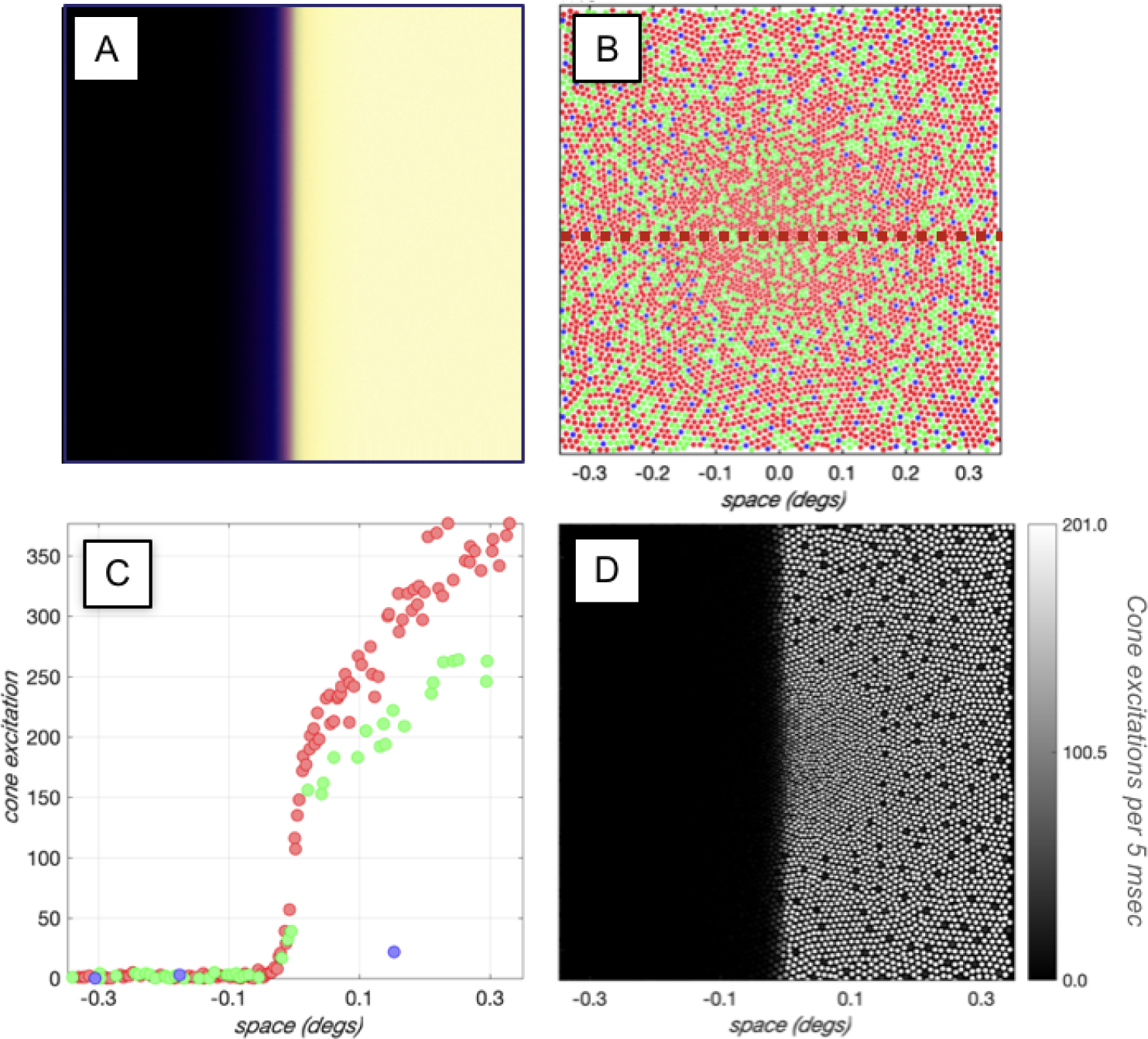
Cone mosaic excitations in response to an edge presented briefly at the fovea. (A) Longitudinal chromatic aberration spreads the short-wavelength light substantially. (B) The cone mosaic samples the retinal irradiance nonuniformly, even in the small region near the central fovea. The differences include cone aperture size, changes in overall sampling density, and changes in the relative sampling density of the three cone types. (C) The number of cone excitations per 5 msec for a line spanning the edge and near the center of the image. The variation in the profile is due to Poisson noise and dark noise (250 spontaneous excitations/cone/second). (D) The number of cone excitations per 5 msec across a small patch near the central fovea. The dark spots are the locations of simulated short-wavelength cones.

It is difficult to have simple intuitions about how these many parameters combine to impact the cone excitations. We hope the ability to perform these computations will help clarify system performance and assist investigators in developing intuitions.

## Discussion

Eye models implemented in the ISET3d and ISETBio software calculate the retinal image and cone excitations of 3D scenes. Knowledge of this encoding may be useful for basic research about depth perception or for applied research into the image quality of novel displays. Accurate calculations of the retinal image from 3D scenes depend strongly on factors we reviewed, including accommodation, pupil diameter, and chromatic aberration. Accounting for these factors is a necessary starting point for building computational models of depth perception and metrics for image quality of 3D displays.

### Ray-tracing is accurate at depth occlusions

The ray-tracing methods we describe accurately capture many features of image formation, but they are compute intensive. In some cases the retinal irradiance can be calculated using simpler methods. It is worth considering when ray-tracing methods are required and when simpler calculations suffice.

The simplest alternative to full ray tracing is to calculate the retinal irradiance by adding the light contributed from each visible scene point to the retinal irradiance (Barsky, Bargteil, Garcia, & Klein, 2002). In this approximation, the light from each scene point is spread in the retinal according to a point spread function; the point spread depends on the distance between the scene point and the focal plane. For some calculations this approach may suffice. An example of such a scene is a large planar surface, such as a flat panel display.

In scenes that include nearby occluding objects this approach can be inaccurate. For example, consider a scene comprised of two planes separated in depth, with the optics in focus for the near plane (Figure 12). We compare retinal images that are computed by convolving each plane separately with the appropriate depth-dependent point spread and summing (panel A), or by ray-tracing (panel B). The two results differs significantly near the occlusion (panel C).

**Figure 12:**
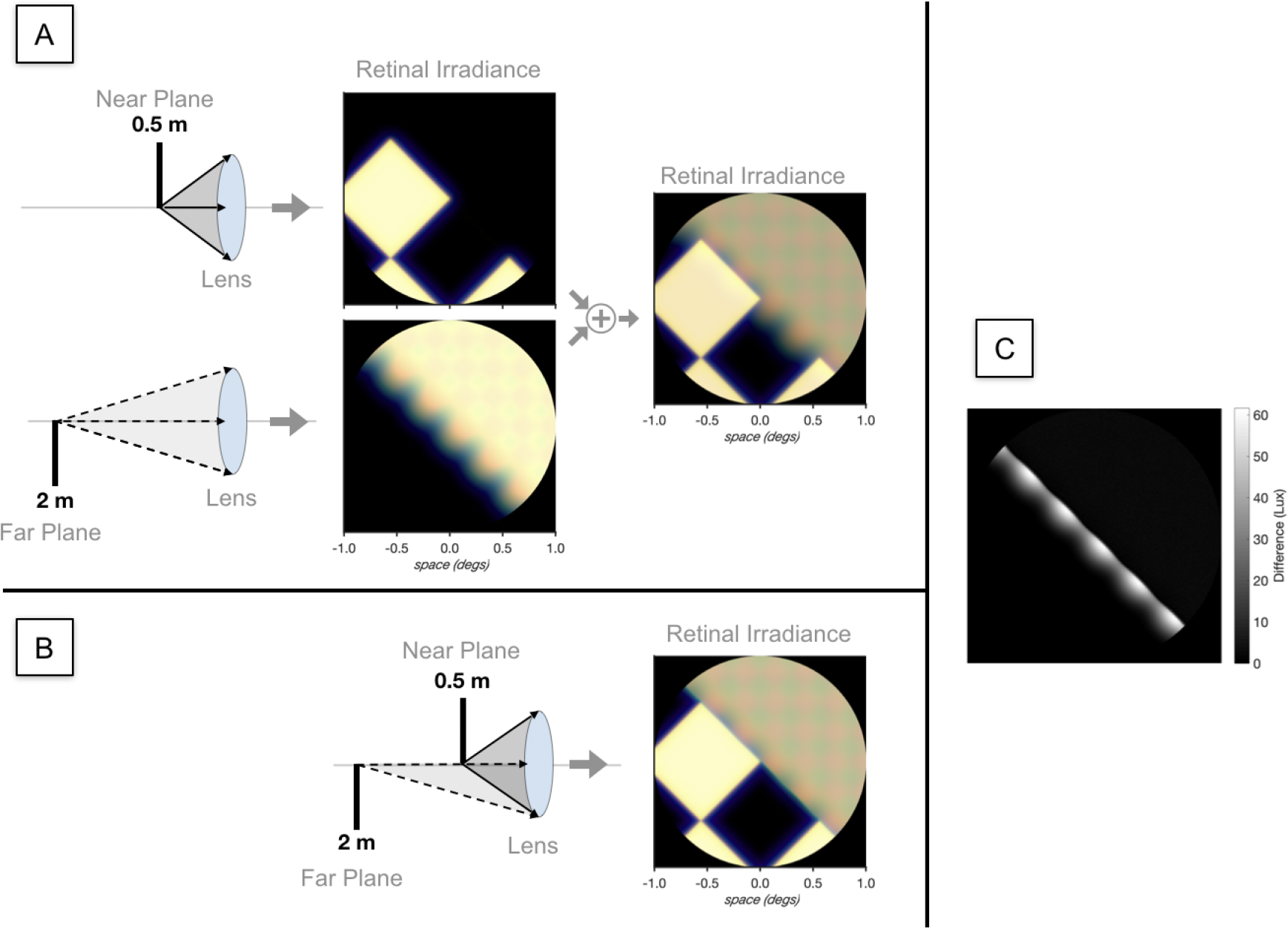
Retinal image calculations near occluding boundaries. The scene comprises two planes with identical checkerboard patterns, placed at 0.5 meters and 2 meters from the eye. The eye is accommodated to the near plane. (A) The two planes are imaged separately, using the correct point spread for each depth. The lens diagrams show the rays for each plane before summation. The irradiance data are then summed. (B) The two planes are rendered using ISET3d ray-tracing. Note that some rays from the far plane are occluded by the near plane. The rendered retinal irradiance is shown (C) The A-B monochromatic difference image (absolute irradiance difference, summed across wavelengths) is large near the occluding boundary. The difference arises because rays are occluded rays from the distant plane.

The difference can be understood by considering the rays arriving at the lens from points on the distant plane near the depth discontinuity. A fraction of these rays are occluded by the near plane, changing the amplitude and position of the point spread function from these points.

The depth-dependent point spread calculation can be approximated in some cases, making it of interest in consumer photography and computer graphics applications (Barsky, Tobias, Chu, & Horn, 2005; Kraus & Strengert, 2007). But the calculations is not always physically accurate because the precise calculation depends on many factors including the position of the occluding objects, the eye position, viewing direction, pupil diameter, and accommodation. To the extent that physically accurate information at depth boundaries is important for the question under study, ray-tracing is preferred.

### Single subject measurements

Schematic eyes typically represent an average performance from a population of observers. It is possible to personalize the surface parameters of a schematic eye for a single subject from adaptive optics measurements of the point spread functions over a range of field heights and angles (Navarro, González, & Hernández-Matamoros, 2006). Using optimization methods, the lens thickness or biconicity of the cornea, can be adjusted so that the model eye matches the point spread function measured in a single subject or for a standard subject (Polans et al., 2015). In this way, an eye model that reflects the properties of an individual subject can be created to estimate a personalized retinal image for three-dimensional scenes.

### Limitations and related work

We continue to expand the range of optics parameters included in the ISET3d eye models. The main extensions are to help clarify effects that arise from normal variation in the eye and extensions necessary to extend the range of model eyes. We are currently adding and testing off-axis and tilted lens positions to understand the impact of decentering. We see value in adding the ability to model gradient-index surfaces, and also to model a larger set of surface shapes, beyond the current group of spherical and biconic. One of the questions we consider is whether the extended modeling is likely to have an impact on the spatial mosaic of cone excitations.

Other investigators have also used ray-tracing to clarify the properties of human image formation, for example in the context of designing intraocular lenses and considering gradient-index lenses (Einighammer, Oltrup, Bende, & Jean, 2009; Schedin, Hallberg, & Behndig, 2016; Gómez-Correa, Coello, Garza-Rivera, Puente, & Chávez-Cerda, 2016). Those papers describe ideas based on ray-tracing principles, and they illustrate their calculations. A number of authors have also described software for using model eyes to calculate the impact of human optics on 3D scenes (Wei, Patkar, & Pai, 2014; Wu, Zheng, Hu, & Xu, 2011; Mostafawy, Kermani, & Lubatschowski, 1997). However, we have not found papers that provide open-source software tools that account for diffraction, spectral properties, and integrate the estimated retinal irradiance with cone mosaic calculations as we do in our tools. The ideas in the literature do provide valuable additional optics calculations that could be incorporated into the open-source and freely available software we provide (https://github.com/iset3d/wiki).

A key limitation of ray-tracing methods is computation time. During development and testing we render small, low resolution images that take roughly a minute to render. A final, high quality 800×800 resolution retinal image can take over an hour to render on an 8-core machine. We are using cloud-scale computing, sending multiple Docker containers in parallel to the cloud, to reduce the total rendering time. We used an associated toolbox, ISETCloud, to implement this parallel rendering for several calculations in this paper.

Finally, we are committed to linking the ISET3d calculations with the ISETBio software (Farrell et al., 2014; Brainard et al., 2015; Cottaris, Jiang, et al., 2018). That software enables users to convert the retinal irradiance into the spatial pattern of excitations of the cone mosaic and to develop models of the neural circuitry response. We hope to expand the combination of tools to understand the effect of different scene radiances in the retina and brain, and to use these analyses to learn more about visual perception and to create tools for visual engineering.

## Acknowledgements

We thank Jennifer Maxwell for her work in developing and documenting the software. This research was supported by a donation from Facebook Reality Labs to Stanfords Center for Image Systems Engineering and to the University of Pennsylvania.

## Appendix

These tables show the parameters of three eye models described in the text.

**Table 1:** Navarro Eye Model (Escudero-Sanz & Navarro, 1999)

| Surface | Radius (mm) | Ocular Media<br>(to next surface) | Conic Constant | Thickness (mm)<br>(to next surface) |
| --- | --- | --- | --- | --- |
| Cornea Anterior | 7.72 | Cornea | -0.26 | 0.55 |
| Cornea Posterior | 6.50 | Aqueous | 0 | 3.05 |
| Lens Anterior | 10.20 | Lens | -3.1316 | 4.00 |
| Lens Posterior | -6.00 | Vitreous | -1.0 | 16.3203 |
| Retina | -12 |  |  |  |
| Refractive Indices |  | (Escudero-Sanz & Navarro, 1999) |  |  |

**Table 2:** Arizona Eye Model (Schwiegerling, 2004)

| Surface | Radius (mm) | Ocular Media<br>(to next surface) | Conic Constant | Thickness (mm)<br>(to next surface) |
| --- | --- | --- | --- | --- |
| Cornea Anterior | 7.8 | Cornea | -0.25 | 0.55 |
| Cornea Posterior | 6.50 | Aqueous | -0.25 | 2.97 |
| Lens Anterior | 12.0 | Lens | -7.518749 | 3.767 |
| Lens Posterior | -5.224557 | Vitreous | -1.353971 | 16.713 |
| Retina | -13.4 |  |  |  |
| Refractive Indices |  | (Schwiegerling, 2004) |  |  |

**Table 3:** Le Grand Full Theoretical Eye (Artal, 2017)

| Surface | Radius (mm) | Ocular Media<br>(to next surface) | Conic Constant | Thickness (mm)<br>(to next surface) |
| --- | --- | --- | --- | --- |
| Cornea Anterior | 7.8 | Cornea | 0 | 0.55 |
| Cornea Posterior | 6.50 | Aqueous | 0 | 3.05 |
| Lens Anterior | 10.20 | Lens | 0 | 4.00 |
| Lens Posterior | -6.00 | Vitreous | 0 | 16.5966 |
| Retina | -13.4 |  |  |  |
| Refractive Indices |  | (Atchison & Smith, 2005) |  |  |

## Footnotes

1 Docker packages software with its dependencies into a container that can run on most platforms. See www.docker.com.

